# Deep-rooted plant species recruit distinct bacterial communities in 3 m deep subsoil

**DOI:** 10.1101/2021.06.02.446747

**Authors:** Frederik Bak, Annemette Lyhne-Kjærbye, Stacie Tardif, Dorte Bodin Dresbøll, Mette H. Nicolaisen

## Abstract

Deep-rooted plants can obtain water and nutrients from the subsurface, making them more resilient to climatic changes such as drought. In addition, the deeper root network also allow the plants to recruit bacteria from a larger reservoir in the soil. These bacteria might contribute to nutrient acquisition and provide other plant beneficial traits to the plant. However, the deep rhizosphere communities’ compositions and their assembly dynamics are unknown. Here, we show, using three perennial crops, Kernza, lucerne and rosinweed, grown in 4 m RootTowers, that deep rhizosphere bacterial communities are plant specific, but clearly distinct from the shallow communities. We found that the diversity decreased with depth in the rhizosphere, whereas abundance of 16S rRNA gene copies did not change with depth in lucerne and rosinweed. Furthermore, we identified a subgroup (4-8%) of ASVs in the rhizosphere communities that could not be retrieved in the corresponding bulk soil communities. The abundances of genes determined by qPCR involved in N-cycling: *amoA, nifH, nirK, nirS* and *nosZ* differed significantly between plant species, suggesting differences in N content in the root exudates of the plant species. Our results suggest that colonization of the rhizosphere by bulk soil bacteria is not limited by carbon supply, but rather by dispersal. Furthermore, the abundance of N cycling genes indicate that deep rhizosphere bacteria have the potential to provide N through nitrogen fixation.

## Introduction

Due to the changing climate, drought spells are expected to increase in length and frequency, thereby threatening agriculture by reducing crop yield (Spinoni, Naumann and Vogt 2017). Furthermore, current agricultural practices have reduced the quality of top soils leading to lower nutrient availability and decreased microbial diversity in many areas (Lupwayi, Rice and Clayton 1998). To reduce dependence on the top soil for nutrient and water supply, deep-rooted cropping systems have been suggested towards a sustainable intensification of crop production (Thorup-Kristensen *et al.* 2020). These systems are expected to show higher resilience towards perturbations in climatic conditions as they can obtain water and nutrients, as well as recruit microorganisms, from the subsurface (Maeght, Rewald and Pierret 2013).

Microorganisms inhabit all parts of the plants and develop complex and dynamic interactions with their host. In the rhizosphere, microorganisms play important roles and have the potential to influence plant health, development and productivity through direct and indirect mechanisms (Berendsen, Pieterse and Bakker 2012). The most abundant microorganisms in the rhizosphere are the bacteria, which are involved in plant beneficial functions such as nutrient acquisition, antagonism against pest and pathogens, as well as activation of plant host defenses against plant stress (Lugtenberg and Kamilova 2009).

In plants with shallow roots, the rhizosphere communities are specific to individual species or even cultivars (Berendsen, Pieterse and Bakker 2012; Bulgarelli *et al.* 2013). This specificity might depend on differences in exudate compositions (Sasse, Martinoia and Northen 2018), although this has been argued to be too simple (Middleton *et al.* 2021). Furthermore, vertical transmission from the seeds have been suggested to be involved in shaping the microbiome of seedlings, even though the main effect is expected at the phyllosphere (Shade, Jacques and Barret 2017). In any case, the current knowledge on rhizosphere bacterial communities of crops are limited to the top soil, with a few studies focusing on the bacterial communities developing in the rhizosphere down to 0.75 m below ground surface (bgs) (Uksa *et al.* 2014). This knowledge gap can partly be explained by the difficulties in accessing the deep roots for sampling (Maeght, Rewald and Pierret 2013).

In the bulk soil, it has been repeatedly shown that bacterial abundance and diversity decrease with depth (Fierer, Schimel and Holden 2003; Eilers *et al.* 2012; Bak *et al.* 2019), coinciding with a decrease in organic carbon and nutrients. Not surprisingly, this leads to an increase in the relative abundances of autotrophic and lithotrophic bacteria concurrent with a decline in taxa able to degrade plant polymers (Bak *et al.* 2019). It remains unclear, whether this change in the bulk soil communities with depth will similarly affect the composition of the bacterial communities in the deep rhizosphere. Possibly, the deep bulk soil communities represent a reservoir for recruitment of bacterial rhizosphere communities, and their lower abundance and diversity could lead to reduced diversity of the deep rhizosphere communities. Despite this, the deep rhizosphere communities might depend heavily on root exudates for nutrition. Consequently, bacterial communities of the deep soil rhizosphere communities might be less similar between different plant species due to differences in exudate composition. Hence, more knowledge on the bacterial communities in the deep rhizosphere, and where the communities are recruited from, will be important for developing cropping systems for deep-rooted crops. Additionally, these communities might affect ecosystem processes such as long-term carbon sequestration (Rumpel and Kögel-Knabner 2011) and soil formation.

In the bulk soil, nitrogen content declines down through the soil profile (Hirsh and Weil 2019). For non-legume crops, this would lead to an expected increase in the C:N ratio in the rhizosphere with depth. Furthermore, the need for N in the deep soil layer would be expected to change the drivers for establishment of communities with different functional potential for nitrogen fixation and turn-over in deep rhizosphere communities compared to shallow rhizosphere communities. In a recent study by Piexoto et al. (2020), it was shown that despite a lower rhizodisposition in the lower soil layers, a higher N input into the soil in the deep lucerne rhizosphere resulted in higher C partitioning into the microbial biomass production, and contributed this finding to the N-fixing ability of the root-nodules. Hence, the microbial N-cycle in deep soil layers may impact the C-processes in these soil layers.

The overall aim of this study was to characterize the bacterial communities in the rhizosphere of three perennial crops, lucerne (*Medicago sativa* L. cv. Creno), Kernza (*Thinopyrum intermedium*), and rosinweed (*Silphium integrifolium*), down to 3 m bgs, by asking: 1) do plants recruit rhizosphere communities from the deep bulk soil layers?; 2) are the deep rhizosphere communities more similar than the shallow rhizosphere communities across different plant species?; 3) do the functional potential in N cycling change with depth, and is there an impact of plant species on this potential?

To address these research questions, we grew the crops in the unique RootTower facility, recently developed at University of Copenhagen (Thorup-Kristensen *et al.* 2020). These towers are an important innovation that enable sampling and further studies of deep-rooted cropping systems and the plant-microbe interactions taking place down to 4 m bgs. All three crops develop deep root systems. Kernza has a fibrous root, while rosinweed and the legume lucerne develop taproot systems. We sampled the rhizosphere and the corresponding bulk soil from three depths before and after two years of plant growth. We estimated the bacterial and fungal abundances, as well as bacterial genes involved in N cycling by qPCR and characterized the bacterial communities by 16S rRNA gene amplicon sequencing.

## Materials and Methods

### Experimental setup

The experiments were conducted at the RootTower facility at University of Copenhagen in 4-m deep RootTowers (4 × 1.2 × 0.6 m) (Fig. 1), with each tower divided in two chambers (4 × 1.2 × 0.3) (Peixoto *et al.* 2020; Thorup-Kristensen *et al.* 2020). The towers were filled in May 2016 with three layers of field soil, Topsoil (0-25 cm), and two subsoils; Upper Subsoil (25-200 cm) and Lower Subsoil (200-400 cm). The topsoil was a 50:50 mixture of clayey loam and sandy loam topsoil both from the University of Copenhagen’s experimental farm in Taastrup, Denmark (55°40′ 08.5”N, 12°18′ 19.4”E) (Rasmussen, Thorup-Kristensen and Dresbøll 2020). The clayey loam subsoils were collected from just below the plough layer at an arable field at Store Havelse, Denmark (55°89′ 83.9”N, 12°06′ 52.8”E). All soils were classified as Luvisols according to the World Reference Base for Soil Resources (IUSS Working Group WRB 2015). A soil bulk density of 1.6 g m^−3^ was obtained in the towers, corresponding to bulk density at field conditions (Thorup-Kristensen *et al.* 2020). Soil characteristics are shown in Table 1.

**Figure 1.**
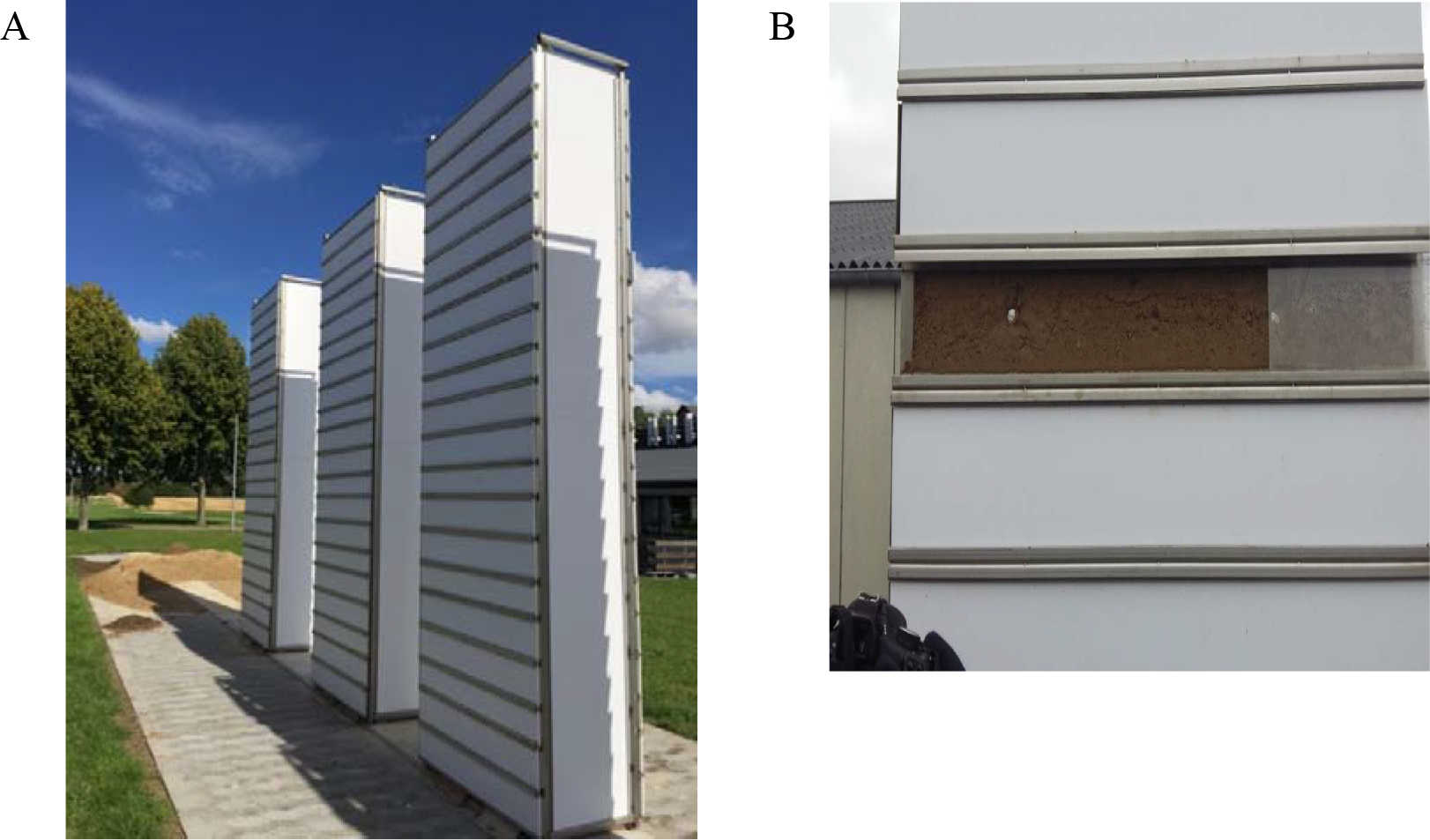
A. Root Towers. B. Access to deep soil layers in the Root Towers.

**Table 1.** Texture of the soil in the RootTowers.

| Depth (cm bgs) | Clay (%)<br><0.002 mm | Silt (%)<br>0.002-0.02 mm | Fine Sand (%)<br>0.02-0.2 mm | Coarse Sand (%)<br>0.2 -2.0 mm | pH<br>(CaCl<sub>2</sub>) | Organic matter (%) |
| --- | --- | --- | --- | --- | --- | --- |
| 0-25 | 8.7 | 8.6 | 46 | 35 | 6.8 | 2.0 |
| 25-200 | 18 | 4.2 | 51 | 27 | 7.4 | 0.3 |
| 200-400 | 16 | 8.7 | 54 | 21 | 7.5 | <0.1 |

The perennial species, lucerne (*Medicago sativa* L. cv. Creno), Kernza (*Thinopyrum intermedium*) and rosinweed (*Silphium integrifolium*) were used for the experiments. Three-months-old Kernza plants and one-year-old lucerne plants from the field, as well as two-months-old rosinweed plants grown in pots with field soil in the greenhouse, were transplanted into individual chambers on 5 July 2016. Plants were selected to have similar root length when transplanted. Each plant was grown in monoculture in three randomized replicates, resulting in nine chambers in total. To mimic the normal plant density in the field, chambers with Kernza and lucerne each contained 5 plants per chamber, all planted with 19 cm equal distance between, while rosinweed contained 3 plants per root chamber, with 28.5 cm between the plants.

### Sampling

The original bulk soil was sampled from the Root Towers before any plants were planted in the summer 2016, and used as reference soil (n = 4 for each depth). Bulk soil and root-associated samples (i.e. roots with attached rhizosphere soil) were collected from the RootTowers during the summer in 2018. Sampling was done for three depths (n = 3): 0-20 cm bgs (topsoil), 160 - 180 cm bgs (upper subsoil), and 280-300 cm bgs (lower subsoil) for each of the nine RootTowers. Hereafter, we refer to the depths as 10 cm bgs (0-20 cm), 170 cm bgs (160-180 cm) and 290 cm bgs (280 – 300 cm). Samples were taken with a sterile soil sampling tube (15 cm long, 1.5 cm in diameter), transferred to sterile Petri dishes, and kept at 4°C until further processing in the lab (done within nine days). Using sterilized tweezers, roots were recovered, shaken to remove excess soil and subsequently transferred into sterile Eppendorf tubes. Hence the original samples were divided into bulk soil and rhizosphere.

### DNA extraction, library preparation and sequencing

All samples were freeze-dried overnight and homogenized by grinding. DNA from approx. 0.25 g of freeze-dried and homogenized sample was extracted using DNeasy Powersoil Kit (Qiagen) following the manufacturer’s protocol. For some of the deep root samples, less material was obtained for extraction. This was accounted for by normalizing to sample mass in the qPCR assay. The DNA concentration and purity were determined using a NanoDrop ND-1000 spectrophotometer (Thermo Fisher Scientific, Carlsbad, Ca, USA) and a Qubit 2.0 fluorometer (Thermo Fisher Scientific, Carlsbad, Ca, USA). Amplicon libraries were prepared by Macrogen Inc. (Seoul, Rep. of Korea) using the primer pair 341F (5’-CCTAYGGGGRBGCASCAG-3’) and 805R (5’-GACTACNNGGGTATCTAAT-3’) (found in Supplementary Table S1) targeting the variable regions V3-V4 of the 16S rRNA gene. Resulting libraries were sequenced on an Illumina MiSeq platform (2 x 300 bp) by Macrogen Inc. (Seoul, Rep. of Korea). Raw sequences will be deposited in the NCBI Sequence Read Archive and are available from the authors upon request.

### Sequence processing

Raw reads were treating using DADA2 version 1.14.1. The protocol for DADA2 was followed using default parameters, with a few modifications. In brief, reads were quality checked and primers were removed using trimLeft in the filterAndTrim() function. The forward and reverse reads were trimmed to 280 and 210 bp, respectively, while the maxEE was set to 3 and 6 for forward and reverse reads, respectively. Detection of amplicon sequence variants (ASVs) were done using the pseudo-pool option. Merged reads in the range of 395 to 439 bp were kept, as reads outside this range are considered too long or too short for the sequenced region. Taxonomy was assigned using the Ribosomal Database Project (RDP) classifier (Wang *et al.* 2007) with the Silva database v.138 (Quast *et al.* 2013). ASVs assigned to Mitochondria or Chloroplast, and ASVs that were not classified at the Phylum level, were removed.

### Diversity estimation and statistical analysis

The 16S rRNA data set was analyzed in R version 4.0.2 (R Core Team 2020) using Phyloseq v. 1.34.0 (McMurdie and Holmes 2013), ampvis2 v. 2.6.6 (Andersen *et al.* 2018) and ggVennDiagram v. 0.5.0 (Gao 2021). The α diversity was estimated using Shannon diversity at the genus level in Divnet v. 0.3.7 (Willis and Martin 2020) with default parameters. To determine whether plant and depth had an effect on the bacterial communities, PERMANOVA using the adonis() function in vegan v. 2.5.7 (Oksanen *et al.* 2020) on a Bray-Curtis dissimilarity matrix made on the ASV table. For visualization nonmetric multidimensional scaling (NMDS) plot were constructed based on the Bray-Curtis dissimilarity matrix. Venn diagrams were made based on all ASVs from all samples belonging to a specific treatment. Testing for differences in bacterial and fungal gene abundances was performed using linear and linear mixed-effect models in the basic and nlme package (Pinheiro *et al.* 2012). Model assumptions were verified, and visual inspection of residual plots did not reveal any obvious deviations from homoscedasticity or normality.

### Quantitative PCR of 16S rRNA gene and ITS

The copy numbers of the 16S rRNA gene, functional genes involved in N-cycling (*nifH, nirS, nirK, nosZ* and bacterial *amoA*) and the Internal Transcribed Spacer 1 (ITS1) region were quantified using quantitative PCR as in Garcia-Lemos et al. (2020). Primer sequences and corresponding annealing temperatures can be found in Supplementary table 1. In brief, PCR reactions were performed in 20 μl reaction mixtures containing: 2 μL of template DNA, 1 μl BSA (20 mg/ml) (New England Biolabs Inc., Ipswich, MA, USA), 10 μl Brilliant III Ultra-Fast SYBR® Green Low ROX qPCR Master Mix (Agilent Technologies, Santa Clara, CA, USA) and 0.8 μl of each primer (10 μM). The qPCR was performed using an AriaMX Real-Time PCR System (Agilent Technologies, Santa Clara, CA, USA). The thermal cycling conditions were 3 min at 95°C followed by 40 cycles of 20 s at 95°C and 30 s at 55-63°C (Supplementary table 1). A final melting curve was included according to the default settings of the AriaMx qPCR software (Agilent Technologies, Santa Clara, CA, USA).

Standard curves for the 16S rRNA gene and the genes involved in N cycling were prepared as in (Garcia-Lemos *et al.* 2020). For ITS1, the standard curve was constructed based on fragments amplified from *Penicillium aculeatum* using the primer pair ITS1F and ITS2 (Supplementary Table 1). Tenfold dilution series were performed for each standard curve. Standard curves spanned a dynamic range from 10^2^ to 10^8^ copies/μL. The reaction efficiencies were between 80 and 106% (see Supplementary Table 1).

## Results

### Bacterial and fungal abundance

To investigate the effects of depth and plant species on the rhizosphere microbial communities of the three deep-rooted plant species, Kernza, lucerne and rosinweed, bulk soil and rhizosphere samples were taken at three depths (10, 170 and 290 cm bgs) from the RootTowers after two years of plant growth. The organic matter decreased in the RootTowers with depth, coinciding with an increase in pH.

The bacterial and fungal abundances were significantly higher in the rhizosphere of the three plant species at all depths compared to the bulk soil (Fig. 2) (p < 0.01). In the rhizosphere, the bacterial abundance changed significantly with depth in Kernza (p =0.02), while there was no significant change in bacterial abundance with depth for lucerne or rosinweed. The fungal abundance decreased significantly with depth for lucerne (p =0.05) and rosinweed (p < 0.001), whereas no decrease in fungal abundance was observed for Kernza (p = 0.1). Except for the bacterial abundance in the bulk soil where Kernza had been grown, a significant decrease with depth in the bacterial and fungal abundances was observed (Fig. 2).

**Figure 2.**
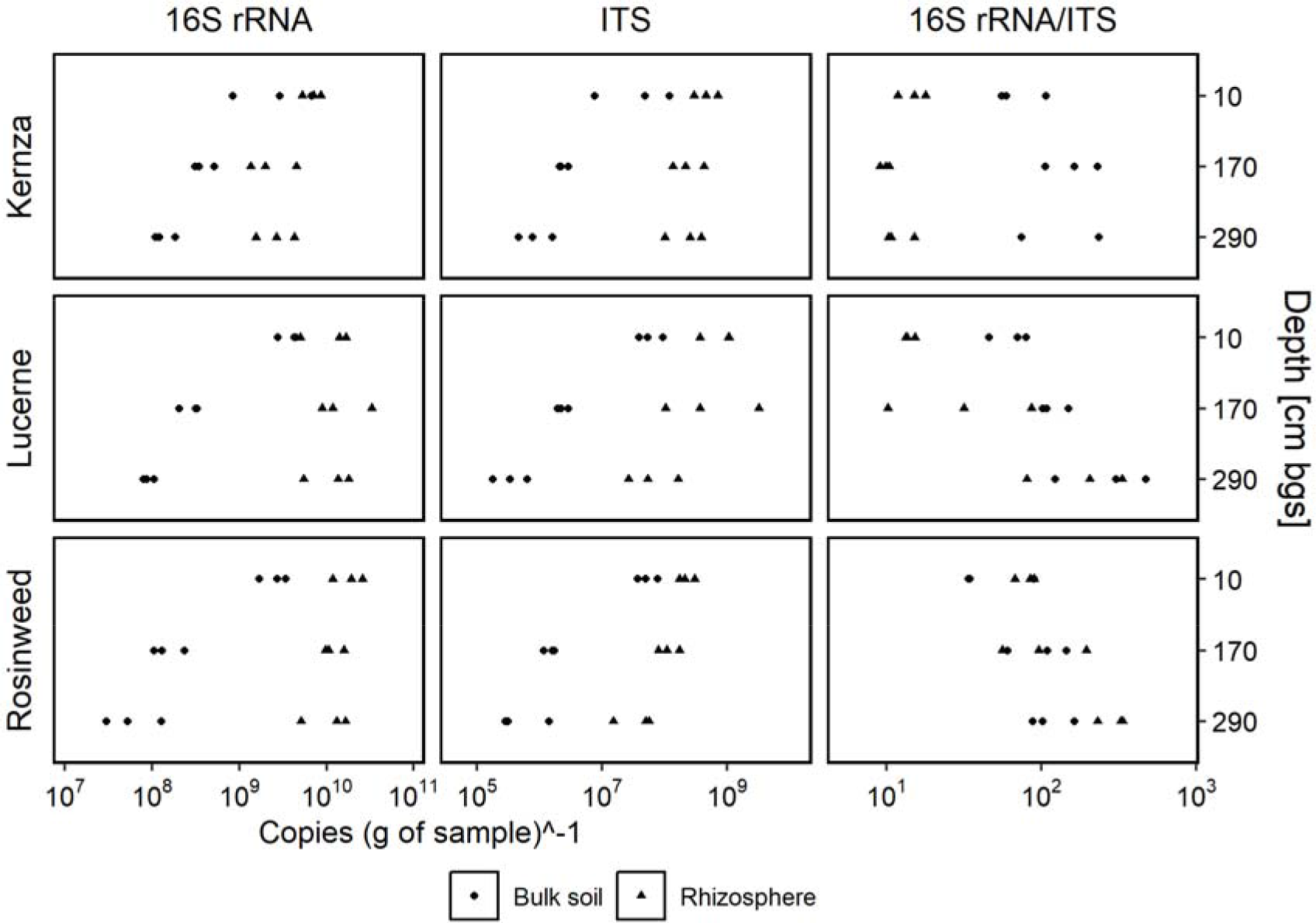
Bacterial and fungal abundances (copies (g of sample)^−1^) based on qPCR of 16S rRNA and ITS region 1, respectively associated with the rhizosphere and bulk soil across three plant species. The ratio between 16S rRNA genes and ITS copy numbers were determined for each sample.

To evaluate the temporal dynamics of the microbial community in the root towers, bulk soil samples taken after two years of plant growth were compared to samples collected before planting. Over the two-year period, there was no change in microbial abundances verifying that the RootTowers did not have an effect on microbial abundance (Supplementary Fig. S1). In addition, plant species did not have any impact on the abundances of 16S rRNA gene or ITS copies in the bulk soil.

In the bulk soil, there was a slight, but significant increase in bacterial:fungal ratio down through the profile, regardless of the plant species that were planted in the tower (Fig. 2). In contrast, significant differences in bacterial:fungal ratios in the rhizosphere were observed between the plant species. In Kernza, the bacterial:fungal ratio did not change with depth in the rhizosphere and was lower than in the bulk soil. In the lucerne rhizosphere, the bacterial:fungal ratio increased with depth (p = 0.03). At 10 cm bgs, the bacterial:fungal ratio was lower in rhizosphere than bulk soil, but there was no difference at 290 cm bgs. For rosinweed, the bacterial:fungal ratio increased with depth (p < 0.001), but was not significantly different from the ratio in the bulk soil at any depth.

### Diversity and Community structure

To obtain more information on the bacterial communities that colonize the deep roots, we performed 16S rRNA gene sequencing of the bacterial community DNA. The total dataset consisted of 66 samples with 4,001,795 reads. The samples contained between 10,281 and 136,479 reads, with a median of 53,480 reads. The dataset comprised 58,720 ASVs. There were no archaeal ASVs in the dataset. The rarefaction curves show that the samples had been sequenced at sufficient depth to capture the richness in the environments, except for the rhizosphere communities in Rosinweed, where a plateau was not reached (Supplementary Fig. S2).

The Shannon diversity of the rhizosphere bacterial communities decreased with depth, with the largest decrease observed for lucerne and rosinweed (Supplementary Fig. S3). Despite this clear trend, significant differences were only observed between the 10 and 290 cm bgs. The Shannon diversity of the bacterial community in the bulk soil was similar at 10 and 170 cm bgs, whereas a significantly lower diversity was observed at 290 cm bgs independent of plant species. At 170 and 290 cm bgs, the Shannon diversity was lower for the rhizosphere compared to the bulk soil for both lucerne and rosinweed, but not Kernza.

NMDS ordination plots based on Bray Curtis dissimilarity showed that the community compositions in the rhizosphere of the three plants were different from those in the bulk soil (Fig 3). In the rhizosphere, we observed a significant effect of depth (PERMANOVA, p < 0.001) and plant species (p < 0.001) (Supplementary Table S2). The interaction between plant and depth was also significant (p < 0.001), indicating a different impact of the plant on the community dependent on the sampling depth. The subsoil rhizosphere communities clustered separately from the top soil communities for all plant species. Additionally, the plot indicated more distinct plant specific communities at 170 and 290 cm bgs compared to 10 cm bgs (Supplementary Table S3). However, the dissimilarity between rhizosphere bacterial communities with-in plant species also increased with depth for the three plant species. A clear separation of the rosinweed rhizosphere community from the rhizosphere communities for Kernza and lucerne was observed.

**Figure 3.**
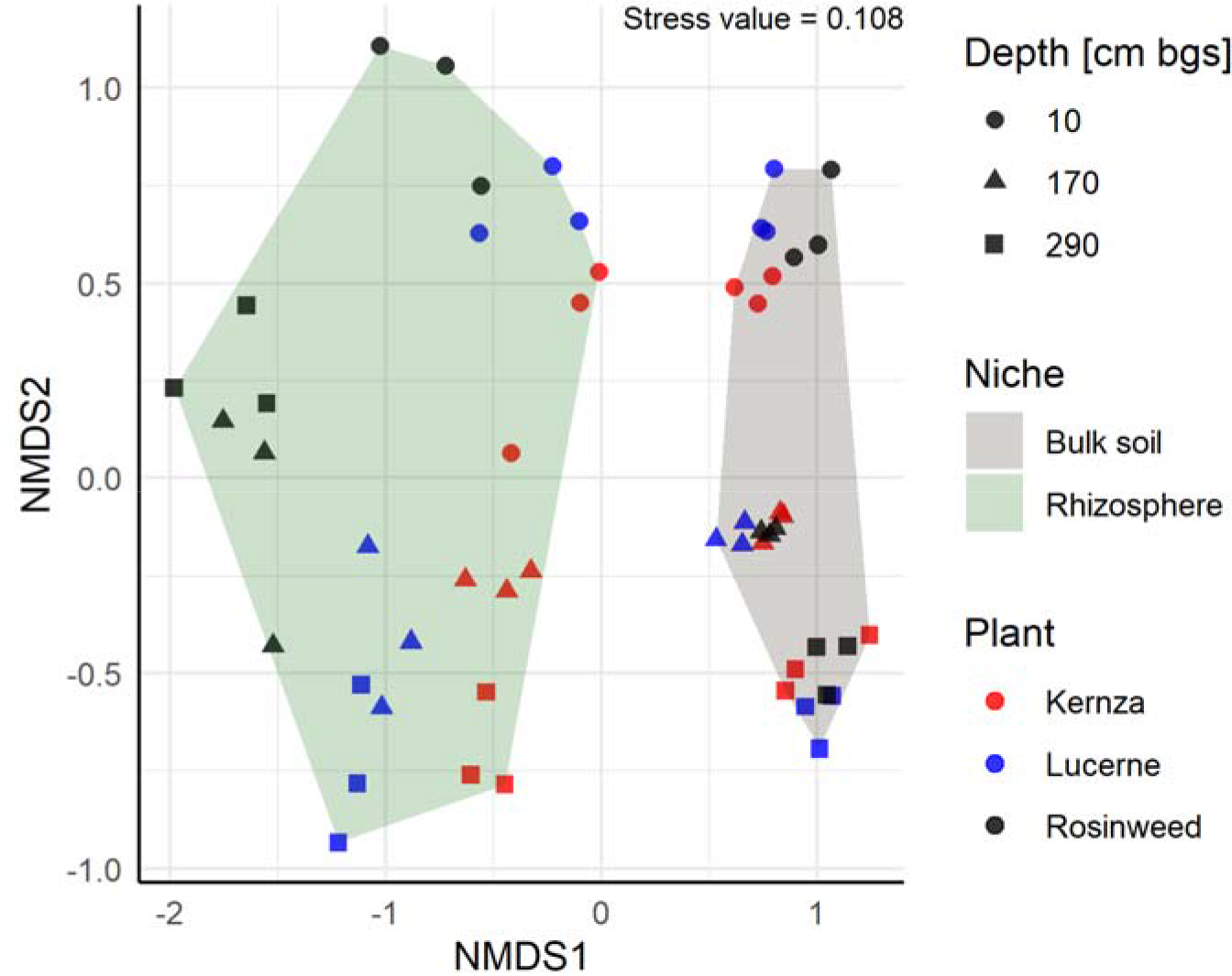
NMDS ordinations of rhizosphere communities and bulk soil communities based on Bray-Curtis dissimilarities.

There were no significant differences in the community compositions between the bulk soil samples coming from the same depth across the three plant species, indicating high reproducibility across towers, and the sampling of true bulk soil in each of the towers (Fig. 3 and Table S2). A change in community structure of the bulk communities was observed between the start and the end of the experiment (Supplementary Fig. S4).

We speculated that the bacterial communities in the deep rhizosphere microbiomes would share more ASVs than the communities in the upper rhizosphere. This is because of the low bacterial diversity in deep bulk soil layers. In contrast to this assumption, the proportion of ASVs shared between plants at different depths, decreased with depth from 19.8% at 10 cm bgs to 12.9% at 290 cm bgs (Supplementary Fig. S5). Furthermore, the proportion of ASVs shared by two plants also decreased with depth.

### Community composition

The rhizosphere communities of all plant species were dominated by genera from four classes, namely Gammaproteobacteria, Alphaproteobacteria, Bacteroidia and Actinobacteria (Supplementary Fig. S6). While bulk soil communities also comprised genera from these four phyla, the diversity was higher and genera belonging to the classes Thermoleophilia, Verrucomicrobiae, Blastocatellia, Bacilli and Acidimicrobia were also among the 15 most abundant genera (Supplmentary Fig. S6D). Interestingly, six bacterial genera (*Pseudomonas, Rhizobium, Streptomyces, Pseudarthrobacter, Niastella* and *Flavobacterium*) accounted for more than 2% of the community in each plant species in at least one depth, suggesting that these genera may be main rhizosphere colonizers (Fig. 4). Generally, these genera increased in relative abundance with depth. The relative abundance of *Pseudomonas* increased with depth for all plant species. However, whereas it constituted above 20% of the communities in rosinweed and lucerne at 290 cm bgs, it constituted less than 5% of the Kernza rhizosphere community. In addition to *Pseudomonas*, the genera *Rhizobium, Ensifer* and *Pseudarhtrobacter* were major constituents of the microbial communities of rosinweed, lucerne and Kernza, respectively (Fig. 4). All these presented genera increased in relative abundance in the deeper rhizosphere.

**Figure 4.**
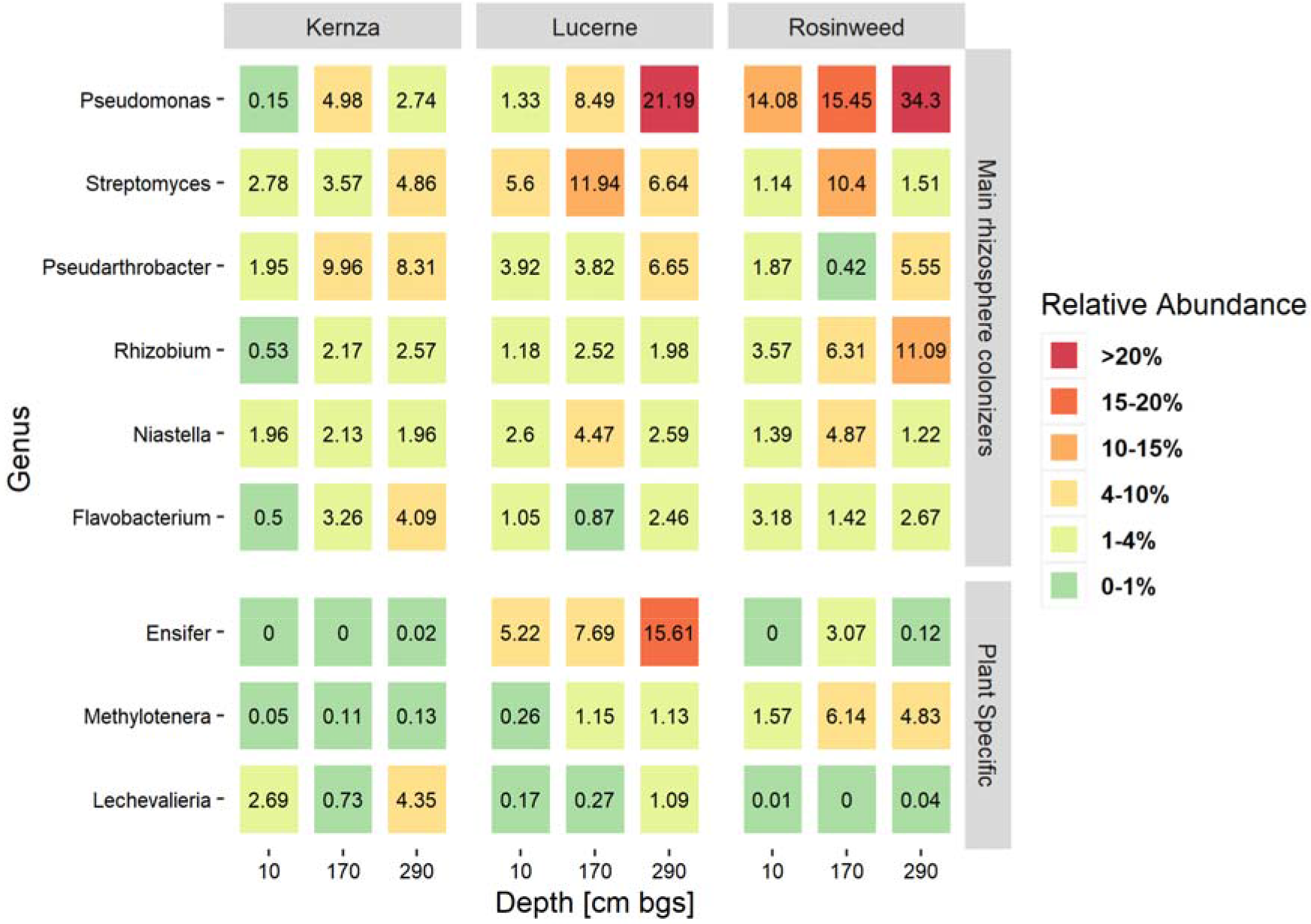
The six most abundant genera across all plant species, classified as main rhizosphere colonizers. Three genera that have particularly high relative abundances in one of the plant species, classified as plant specific genera. Values are in mean relative abundances (n = 3).

Some genera were primarily associated with specific plant genera and found at relative high proportions, but in low abundance in the other plant species (Fig. 4). Deeper roots of rosinweed were colonized by *Methylotenera*, *Ensifer* heavily colonized lucerne roots at all depths, and *Lechevalieria* had a specific association with Kernza.

### Origin of the rhizosphere ASVs

We asked the question whether rhizosphere ASVs would be recruited at the different depths pointing to colonization from the adjacent bulk soil layer (as opposed to vertical transmission from the top-soil). Bulk soil ASVs were categorized as unique to a soil depth or as shared between all soil depths (Supplementary Fig. S7). A total of 6,378 bulk soil ASVs were classified as shared between all soil depths, while the soil at 10 cm bgs contained most unique ASVs (12,809) as compared to 170 cm bgs (10,920) and 290 cm bgs (9,878). In the case of the rhizosphere, the majority of ASVs in the rhizosphere communities of all three plants at the three depths belonged to the shared group of bulk soil ASVs (67-76%) (Fig. 5). Of the rhizosphere ASVs classified as unique to a given depth in the bulk soil, the group belonging to 10 cm bgs constituted the largest proportion (11-14%) of the rhizosphere communities in the three plants at 10 cm bgs, while they accounted for <2% in at 170 and 290 cm bgs. Rhizosphere ASVs unique to 170 or 290 cm bgs in the bulk soil comprised less than 4% of the rhizosphere communities, suggesting a declining importance of recruitment from adjacent bulk soil with increasing depth. Interestingly, for all three plants, a subgroup of ASVs (4-8%) did not belong to any of the bulk soil categories. These ASVs belonged to the genera *Flavobacterium, Pseudarthrobacter, Ohtakweangia, Masillia, Cellvibrio, Duganella, Ensifer* and *Cellvibrio* (Fig. 6).

**Figure 5.**
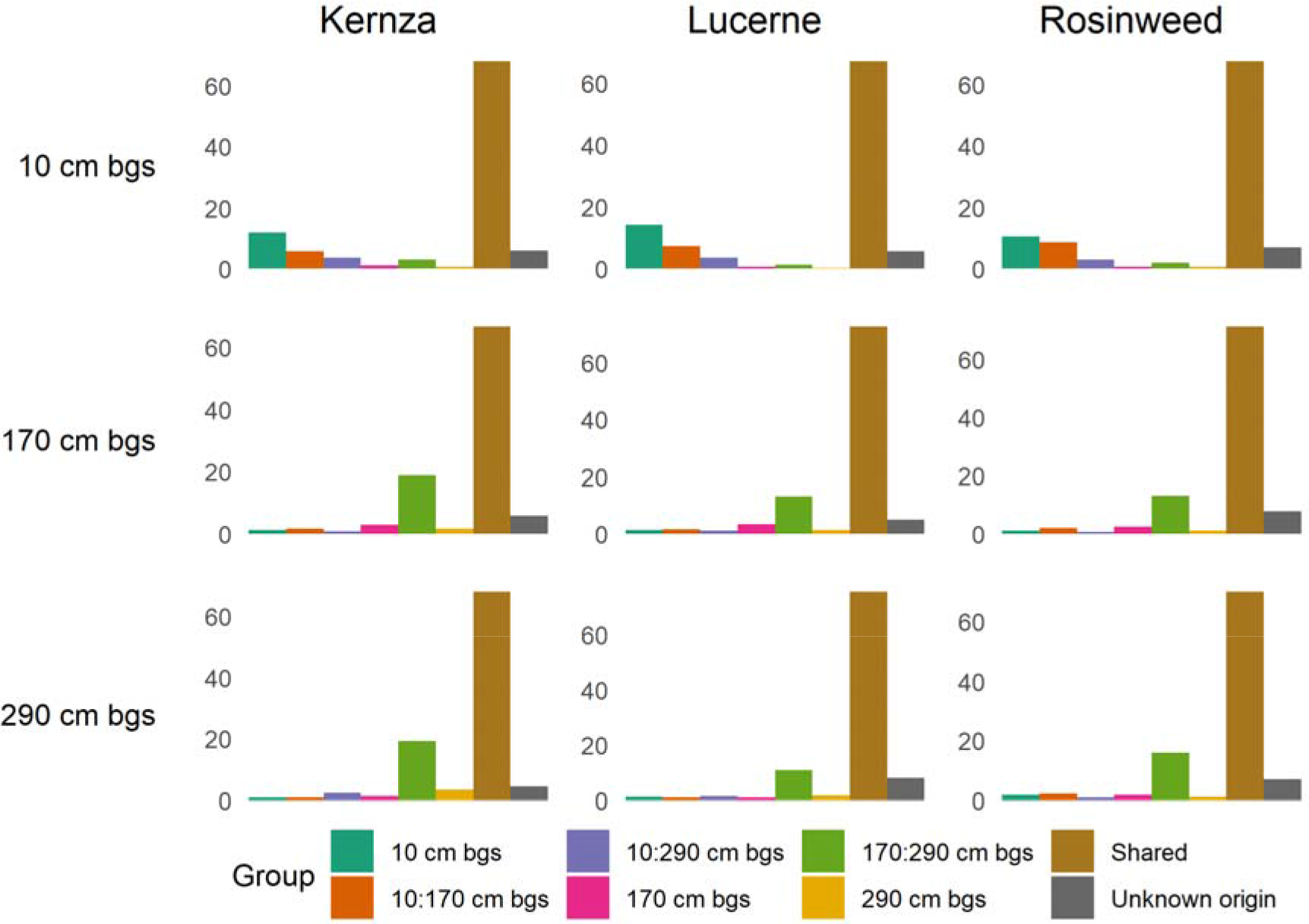
Tracing the potential origin of the rhizosphere ASVs from the bulk soil. Bulk soil ASVs where grouped based on presence/absence at different depths in the rhizosphere. Shared bulk soil ASVs were shared between all three soil depths. Values on the y-axis are relative abundance (%). Unknown origin refers to ASVs that were not found in the bulk soil.

**Figure 6.**
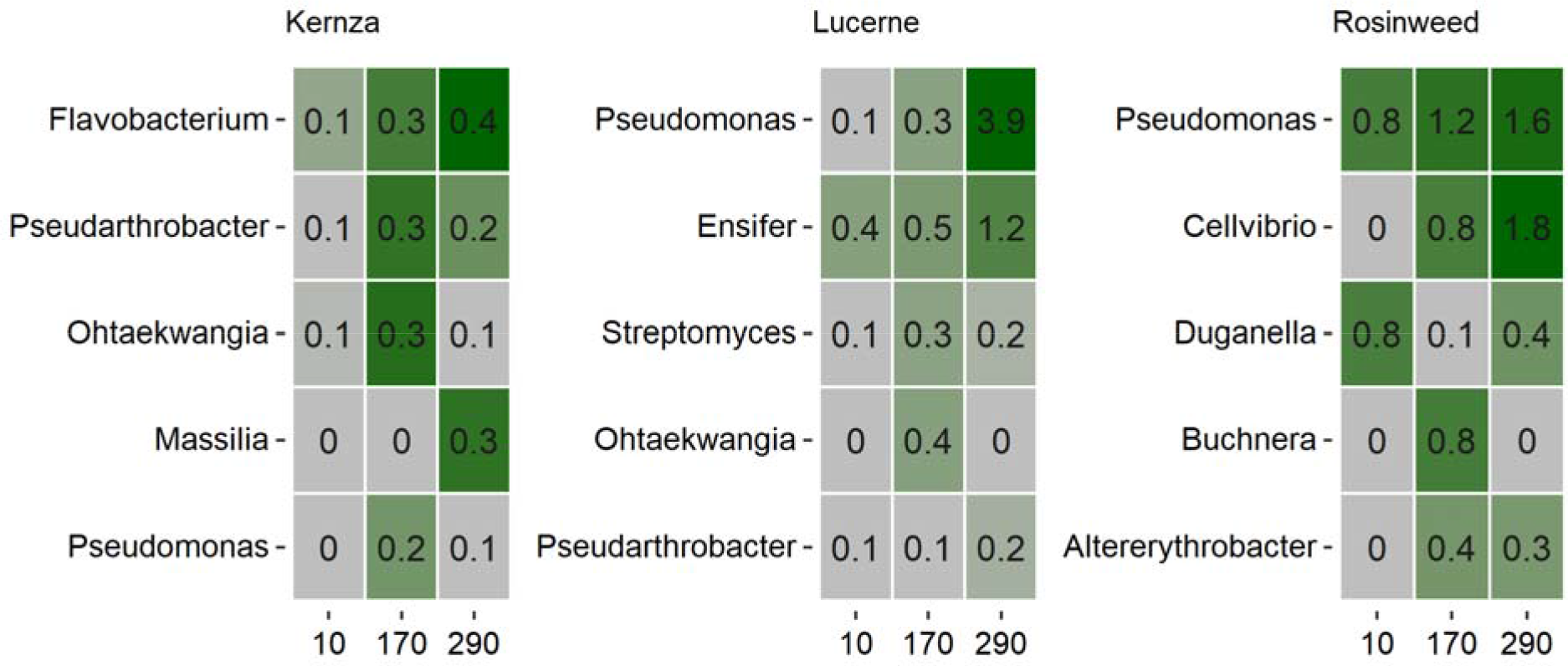
The relative abundance of the five most abundant genera with ASVs that belonged to the group of unknown origin. Numbers on the x-axis indicate depth (cm bgs)

**Figure 7.**
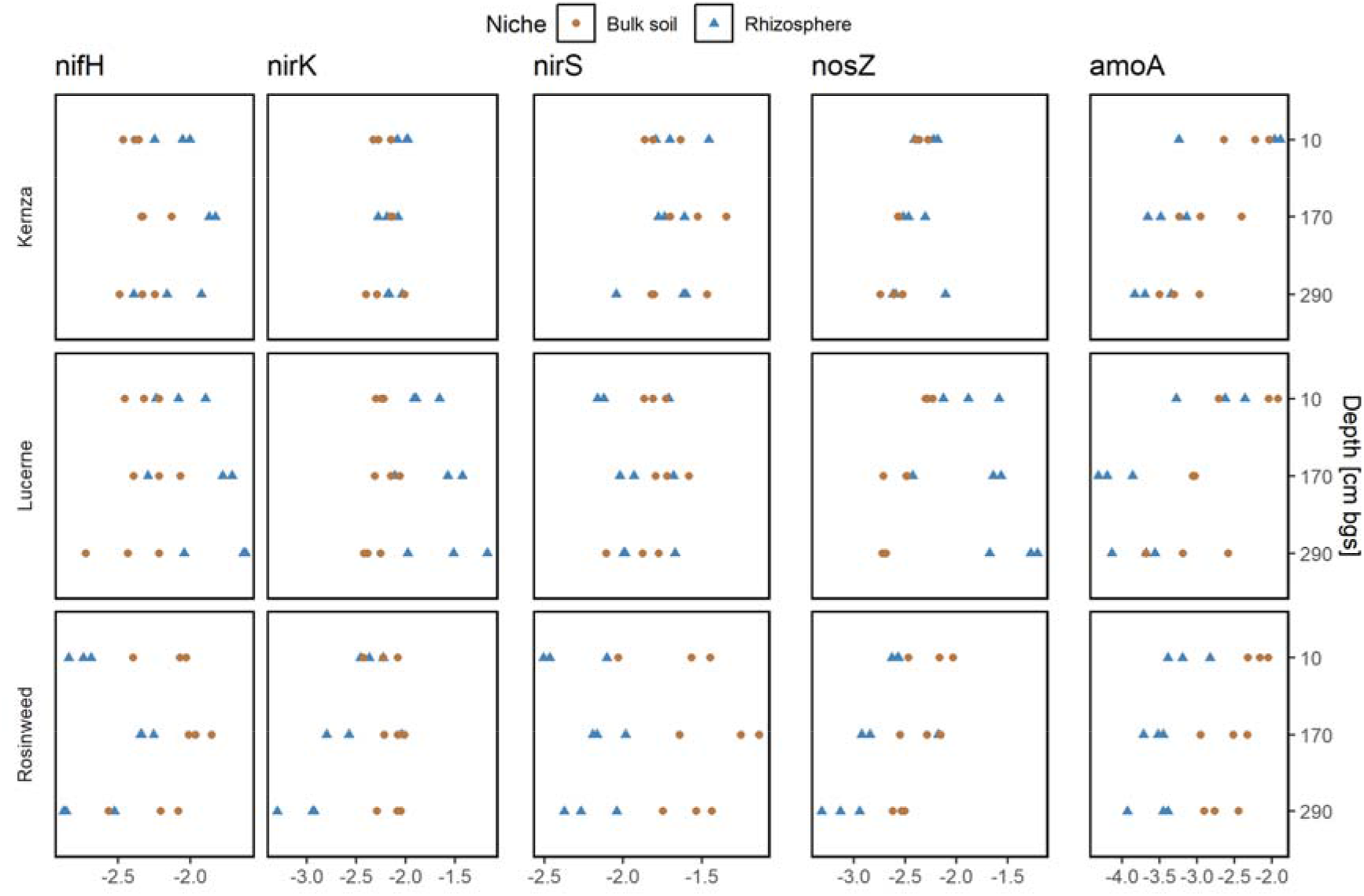
qPCR data of N-cycling genes. The values are standardized per 16S rRNA gene.

### N cycling genes

Quantitative PCR was used to quantify the relative abundance of N-cycling genes at the different depths. The relative abundance of the *nifH* gene involved in N-fixation did not change with depth for Kernza and lucerne; however, both plant species showed a higher relative abundance in the rhizosphere than in the bulk soil, pointing to a recruitment of *nifH* bearing organisms to the roots. Rosinweed had a completely different profile, where the middle depth at 170 cm bgs had an increased relative abundance compared to the other depths. Furthermore, the relative abundance of *nifH* genes was lower in the rhizosphere than in the bulk soil.

The *amoA* gene involved in nitrification showed a general trend of decreased abundance with depth, and furthermore a lower relative abundance in the rhizosphere compared to the bulk soil. For genes *nirS, nirK* and *nosZ* involved in denitrification, there was no difference in relative abundance for Kernza with depth, nor was there any difference between the rhizosphere and the bulk soil for any of the genes. Lucerne had a higher relative abundance of the *nirK* and the *nosZ* genes in the rhizosphere, whereas there was no difference between bulk soil and rhizosphere for the *nirS* gene. The relative abundance did not change with depth, neither in the rhizosphere nor in the bulk soil. In contrast to the other plant species, the Rosinweed rhizosphere had a lower relative abundance than the bulk soil for all three genes at all depths.

## Discussion

In this study, we examined the bacterial communities of three deep-rooted crops. In addition to achieve knowledge on the general community structure at the deep roots, we also analyzed the origin of the bacterial communities as well as the nitrogen cycling potential in the rhizosphere.

### Abundance and rhizosphere effect

Contrary to what we expected, the abundances of bacteria and fungi were stable in the rhizosphere for the three plant species even up to 3 m bgs. It was recently shown that the rhizodeposition decreased significantly with depth, leading us to expect a concurrent decrease in microbial abundance (Peixoto *et al.* 2020). In contrast, the microbial abundance decreased in the corresponding bulk soil, concurrent with a decrease in organic matter in accordance with the literature (Agnelli *et al.* 2004; Eilers *et al.* 2012). Hence, the abundance of microorganisms does not seem to be limited by carbon supply in the root exudates. Instead, access to the root or potential to disperse might be governing factors for microbial populations in the rhizosphere. The general decrease in diversity with depth, especially for lucerne and rosinweed, support this, and is in accordance with the observation of fewer motile strains in the subsoil previously reported (Krüger *et al.* 2019).

The bacterial:fungal ratio (measured as the 16S rRNA gene/ITS gene ratio) increased significantly down through the soil profile. This could be explained by a general decrease in available carbon for heterotrophic growth in deeper soil layers. For the Kernza rhizosphere, the notion of a low bacterial:fungal ratio down through the soil profile, could hint to a close interaction with a fungal community, and maybe even suggest the presence of fungal endophytes in this plant species. However, this will need more research for confirmation.

In accordance with the literature, we found the plant species to shape their rhizosphere bacterial rhizosphere communities in the topsoil (Berendsen, Pieterse and Bakker 2012). Due to lower diversity and abundance of bacteria in the subsoil compared to the topsoil as found in previous studies (Fierer, Schimel and Holden 2003; Eilers *et al.* 2012; Bak *et al.* 2019), we expected that the deep rhizosphere communities would become more similar with depth. Our results, however, showed an increasing effect of plant species on the rhizosphere communities. In addition to a lower diversity in subsoils, the recruitment base (i.e., the bulk soil) is more heterogeneous in subsoils, which could also impact the diversity measure, and explain the higher dissimilarity between replicates from the same plant species. Hence, the similarity between the various plants’ rhizosphere communities decreased with depth. Furthermore, we found a decrease in the proportion of shared ASVs between the plant species at 170 and 290 cm bgs compared to 10 cm bgs. These results imply a stronger selection for bacteria that can colonize the rhizosphere in the subsoil compared to topsoil. This is partly backed up by the decreasing alpha diversity for rosinweed and lucerne. The reason for the stronger selection can be attributed to a decrease in organic carbon with depth, increasing the carbon gradient between the rhizosphere and the bulk soil.

### Composition of bacterial communities

The rhizosphere communities comprised high abundances of *Pseudomonas, Streptomyces, Rhizobium* and *Pseudarthrobacter* across the three plant species, even at 290 cm bgs. These genera contain species well known for their interactions with plants, and expression of plant beneficial traits (Tokala *et al.* 2002; Hayat *et al.* 2010), and especially pseudomonads are well known as colonizers of new habitats when C becomes available (Lugtenberg, Dekkers and Bloemberg 2001), probably due to their copiotrophic lifestyle and r-strategy for growth.

Hence, their dominance in the deep rhizosphere indicates that many beneficial bacterial traits could be available for the deep roots. As inferring functional potential based on taxonomic profiles is difficult, it would be important to verify this by metagenomic sequencing or genome sequencing of isolates.

The high relative abundance of *Rhizobium* and *Ensifer* at 290 cm bgs in rosinweed and lucerne rhizosphere communities, respectively, as well as an increase in abundance of *Rhizobium* in the Kernza rhizosphere with depth, suggests that nitrogen fixation occurs throughout the soil profile, but especially in the deep soil rhizosphere, providing important nitrogen to the plants. Furthermore, it indicates that readily accessible nitrogen for bacteria and plants is limited in the subsurface compared to the topsoil.

With *Ensifer* being a specific colonizer of root nodules of lucerne, it was not surprising to find a high relative abundance of this genus specifically associated with this plant species (Carelli *et al.* 2000). Rosinweed contained a high relative abundance of *Methylotenera*. This genus contains methylotrops that are able to use C1 compounds as sole sources of energy and carbon, especially methylamine (Kalyuzhnaya *et al.* 2006). *Methylotenera* has also been found in the rhizosphere of rice and *Baccaris scandens* (Moronta-Barrios *et al.* 2018; Fuentes *et al.* 2020). The presence of this genus indicates that the root exudates of rosinweed contain a larger fraction of C1 compounds. *Lechevalieria* was found in relative high abundance in the Kernza rhizosphere. While species from this genus has been isolated from rhizosphere in wheat (Zhao *et al.* 2017), no consistent reports have linked this genus to rhizosphere functions.

### Origin of the rhizosphere bacteria

Bacteria in the rhizosphere can originate from the top soil and be transported along the roots or by preferential flow paths (Dibbern *et al.* 2014; Bak *et al.* 2019). Alternatively, they can be recruited from the soil horizons that the roots penetrate. The high abundance of *Pseudomonas* in the deep rhizosphere, coinciding with a drop in diversity with depth, can be explained by dispersal. In a study, on motility in subsurface soil, (Krüger *et al.* 2019) found *Pseudomonas* to be a good disperser in soil samples from 80-120 cm bgs and preferential flow paths from 300-350 cm bgs. Furthremore, motility has been found to be an important trait in initial colonization of sterile roots in a wild type-mutant experiment (Turnbull *et al.* 2001).

The majority of the ASVs identified in the rhizosphere communities could be found in the bulk soil communities. However, a significant portion of approx. 5% of the rhizosphere ASVs were not detected in the bulk soil communities. The genera that comprised these ASVs belong to genera such as *Pseudomonas, Masillia, Niastella, Ohtakweangia, Ferruginibacter* and *Flavobacterium*. Except for *Ferruginibacter*, which has previously only been reported from aquatic habitats, all genera have been found as endophytes in roots or seeds in other crops like durum wheat and maize (Gao *et al.* 2015; Truyens *et al.* 2015; Agnolucci *et al.* 2019). Although we cannot rule out lack of sequencing depth as an explanation, the rarefaction curves obtained for the bulk soil samples indicated that we had sequenced deep enough. Alternatively, the ASVs are pure endophytes, as the entire root was sampled along with the rhizosphere. However, based on previous reports, CFU counts showed 100-fold more bacteria in the rhizosphere compared to the endosphere per gram of sample (Benizri, Baudoin and Guckert 2001; Blain, Helgason and Germida 2017), indicating that not all of the ASVs are endophytes. Hence, we argue that these ASVs originate as seed associated.

### N-cycling potential

For the N-cycling genes, it was found that lucerne had the highest number of denitrifiers, based on quantification of *nirS*, *nirK* and *nosZ* genes. This could be explained by the higher N output from lucerne as compared to the other two species (Peixoto *et al.* 2020). Interestingly, the lucerne rhizosphere seems to recruit denitrifiers not harboring the nirS gene, as these genes were equally abundant in the rhizosphere of lucerne and the bulk soil.

The *nifH* genes were specifically enriched in the lucerne rhizosphere, which would be expected due to the development of root nodules. Interestingly, the nifH genes were also specifically enriched in the Kernza rhizosphere, indicating an increased potential for N-fixation in this habitat. This was in contrast to the rosinweed rhizosphere, where *nifH* genes were not enriched. The ability of recruiting a N-fixing community in the rhizosphere will potentially change the C:N ratio in the Kernza rhizosphere, and thereby impact the not only the N-cycling, but also the incorporation of C into microbial biomass (Peixoto *et al.* 2020).

The current study was performed in field-like RootTowers, providing easy access to the roots, making it possible to sample from the root systems with minimal disturbance of the soil, but it is important to keep in mind that our findings may differ from future findings performed in the field. We did see a change in community structure over time in the bulk soil, however, we did not detect any changes in bacterial or fungal abundances over time. Albeit, that the RootTowers are an artificial system, they pose a promising potential for studying microbe-plant microbe interactions in the depth.

## Conclusion

The three plant species recruited specific rhizosphere communities over the entire sampled region. Furthermore, they have a high proportion of taxa that were shared between bulk soil communities at all depths. Our results suggest that deep-rooted rhizosphere bacterial communities are colonized by species recruited from the surrounding bulk soil, as well as transported down with root growth. However, how important these two mechanisms are in community assembly dynamics remains to be tested in subsequent experiments using quantitative measures. Furthermore, our results pointing towards a substantial proportion of deep rhizosphere communities are seed borne, should be investigated, as this might have implications for the development of microbial inoculants. Deep-rooted crops are receiving increased interest in carbon sequestration. Hence, understanding how communities are shaped can provide practical and applicable knowledge for rhizosphere engineering of these important crops.

## Acknowledgements

We thank J. A. Teem, D.T. Ganzhorn, and M. Schiller S (University of Copenhagen) for their assistance during sampling and D. T. Ganzhorn for the contribution to the qPCR setup. This work was performed as part of the Deep Frontier project, which is a research collaboration between University of Copenhagen, Aarhus University and International Centre for Research in Organic Food Systems (ICROFS), financed by the Villum Foundation.

