## Supplementary information for "Deep-rooted plant species recruit distinct bacterial communities in 3 m deep subsoil"

|  | Page |
| --- | --- |
| Figure S1. Abundance of 16S rRNA gene and ITS region 1 copies | 2 |
| Figure S2. Rarefaction curves 16S rRNA gene reads | 3 |
| Figure S3. Shannon Diversity | 4 |
| Figure S4. NMDS | 5 |
| Figure S5. Venn Diagram | 6 |
| Figure S6. Heatmap of 15 most abundant genera | 7 |
| Figure S7. Venn diagram | 9 |
| Table S1. Primer sequences | 10 |
| Table S2. PERMANOVA results | 11 |
| Table S3. Bray-Curtis dissimilarity | 11 |
| References | 12 |

**Supplemental Figure S1.** Abundance of 16S rRNA genes, ITS region 1 copies and the ratio between the two in bulk soil samples prior to sowing.
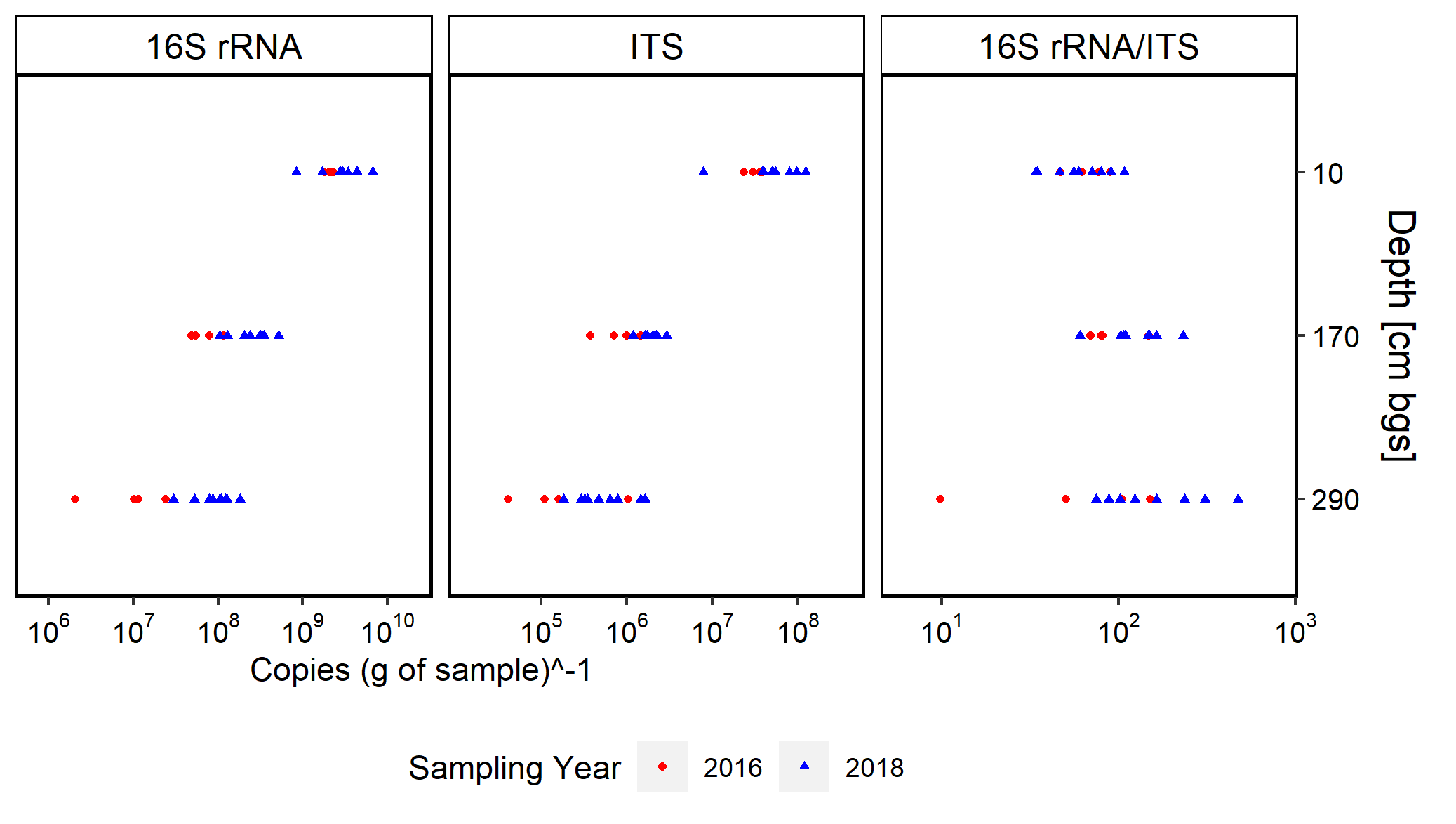

**Supplemental Figure S2.** Rarefaction curves of 16S rRNA gene reads for the different compartments. Sampling depths are indicated at the bottom (cm below ground surface). Bulk Soil samples are from the beginning of the experiment.

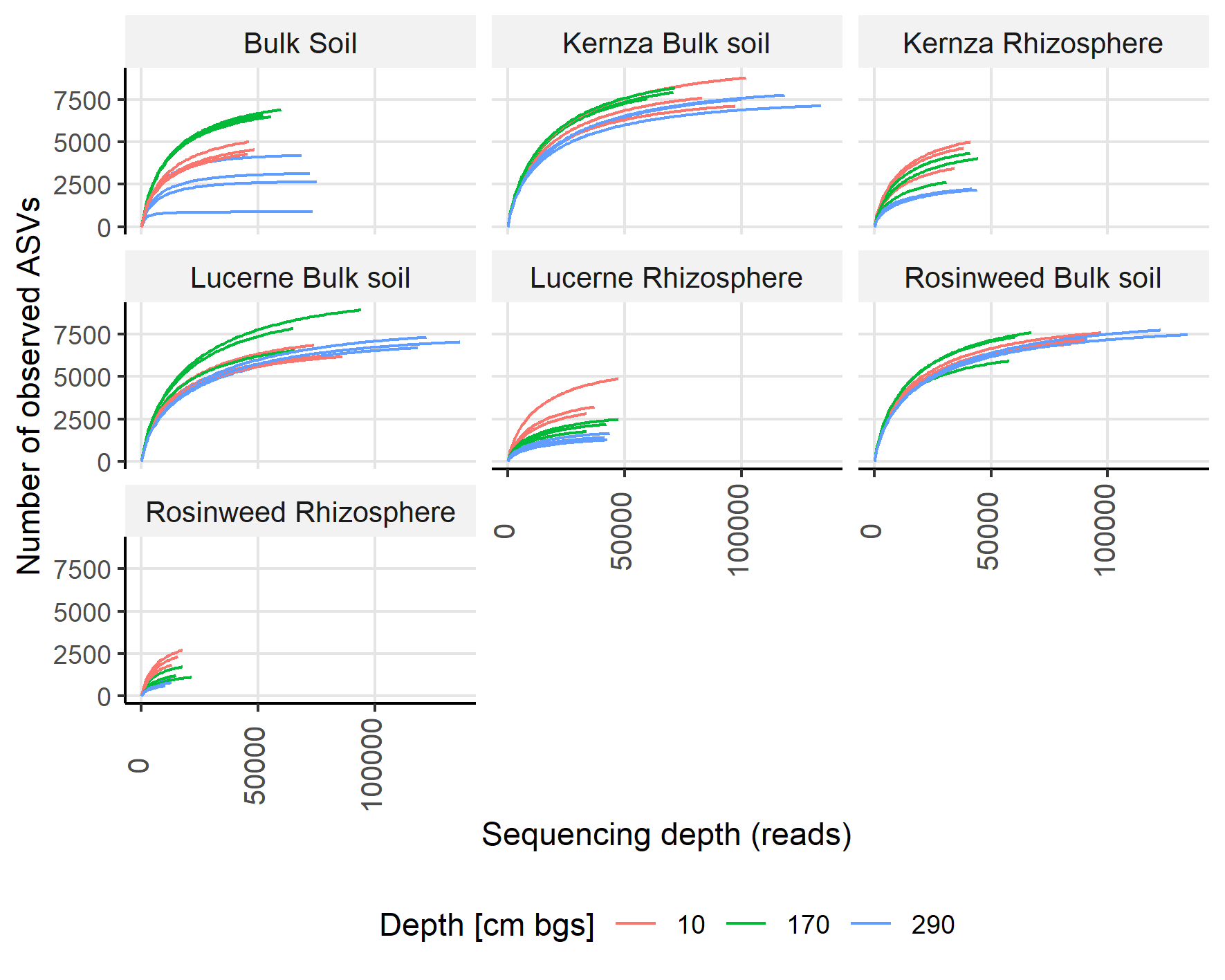

**Supplemental Figure S3**. Shannon Diversity estimated at the genus level of the rhizosphere and bulk soil communities. The mean values and error bars are estimated using Divnet.

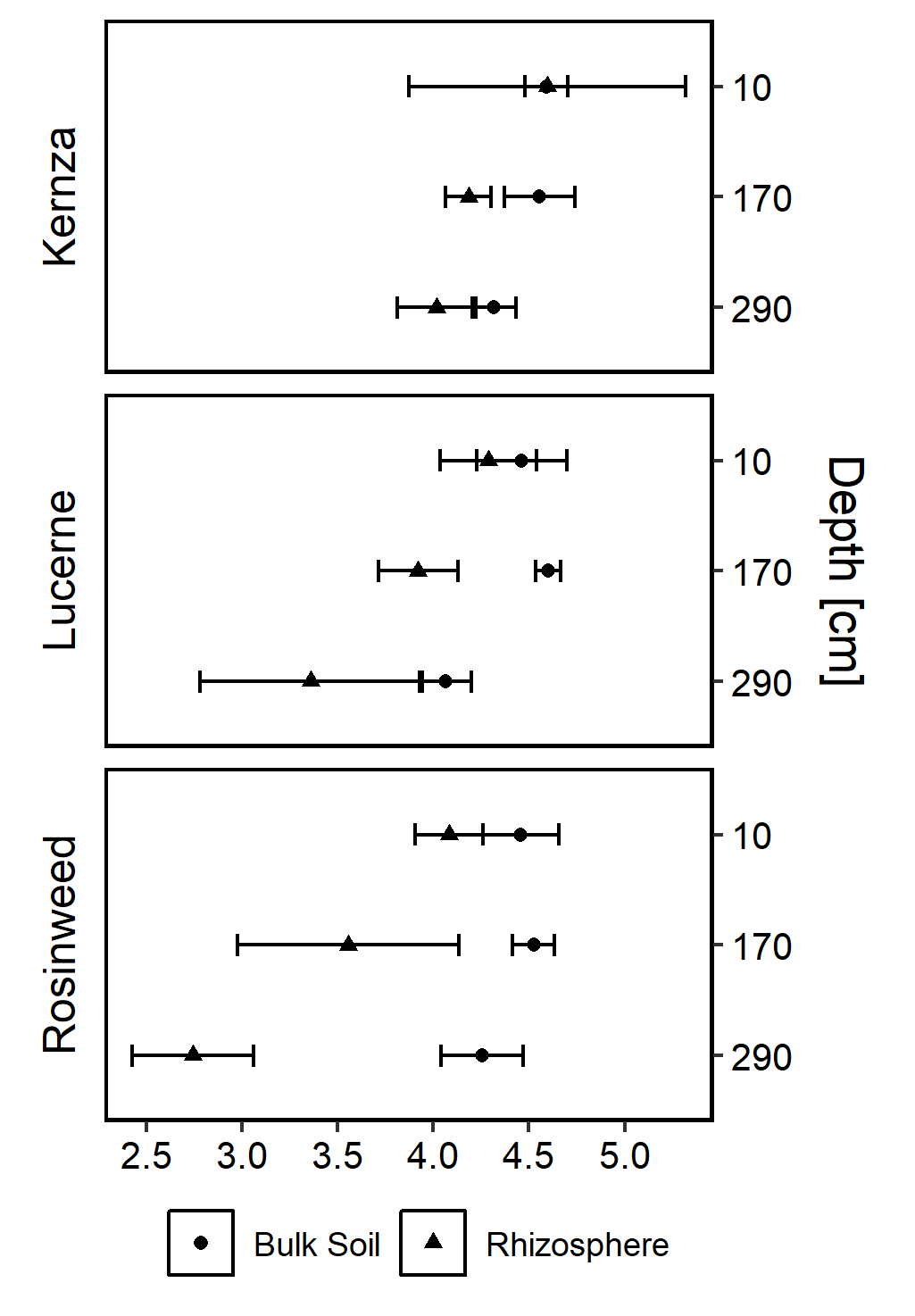

**Supplemental Figure S4.** NMDS ordinations of bulk soil communities based on Bray-Curtis dissimilarities. Bulk Soil samples are from 2016 and 2018.

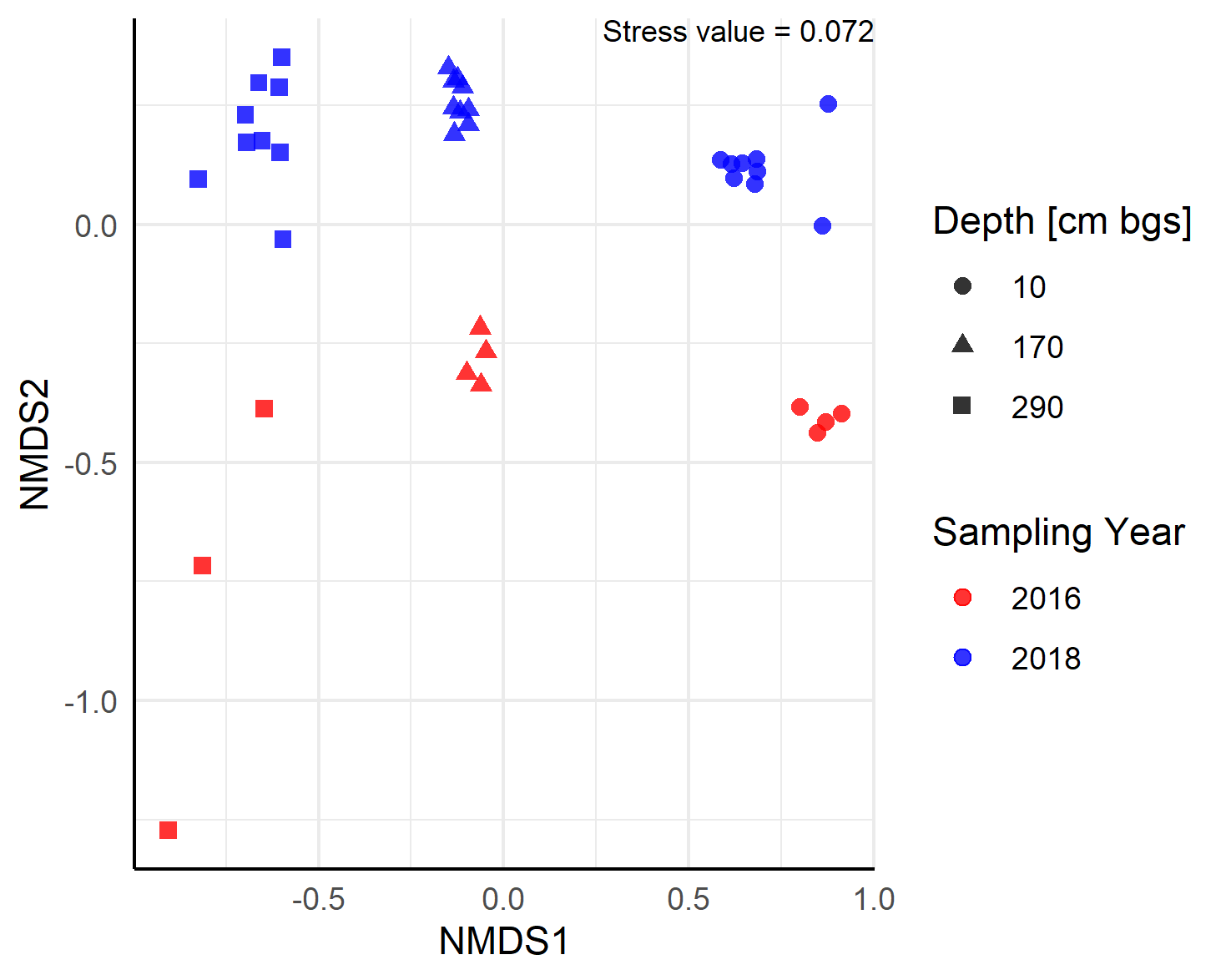

**Supplemental Figure S5.** Venn diagrams of ASVs belonging to the rhizosphere of the Topsoil, Upper Subsoil or Lower Subsoil of the three plants. Values are the proportions of total amount of ASVs that were found in each of the depths, respectively.

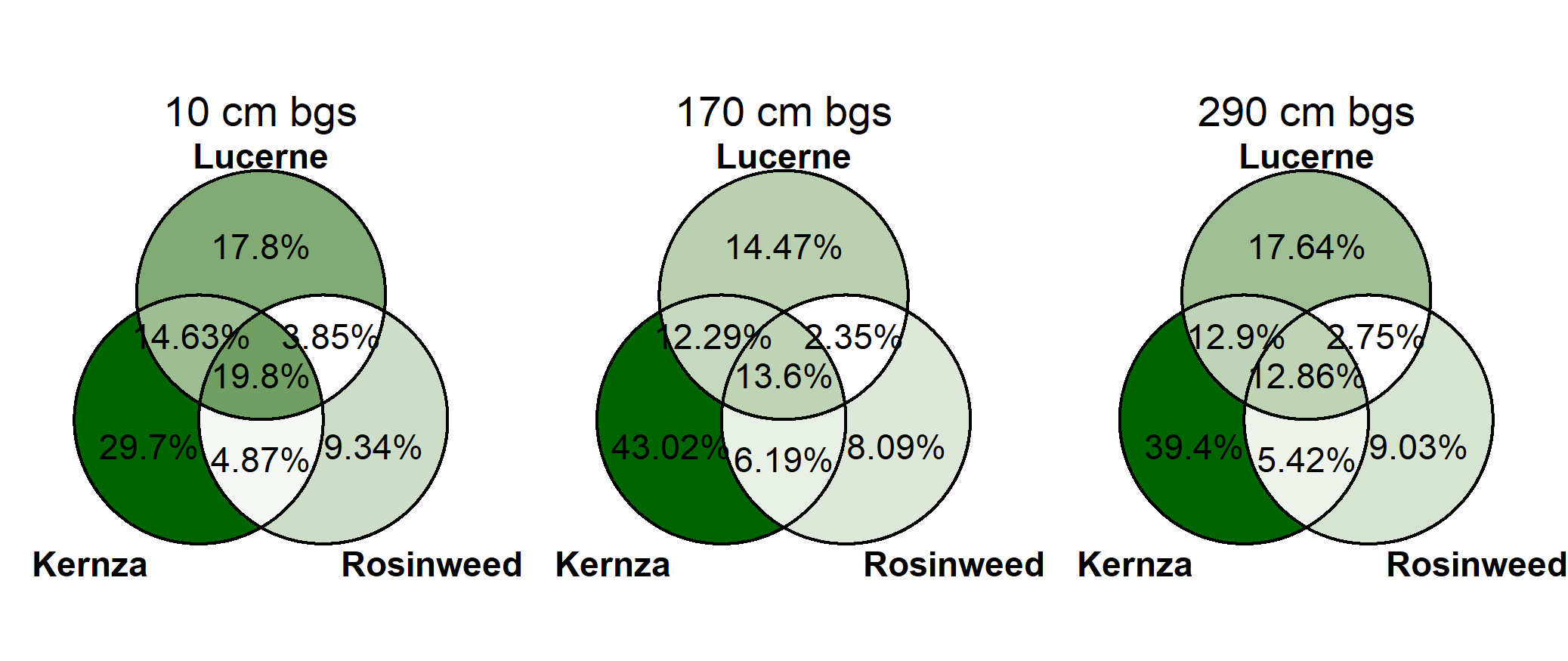

**Supplemental Figure S6.** Heatmap of the 15 most abundant genera across samples for Kernza (A), lucerne (B), rosinweed (C) and bulk soil (D) at the three depths (10, 170 and 290 cm). Values are relative abundance (%). Relative abundances for each replicate are shown for the rhizosphere as high variation between replicates were observed.

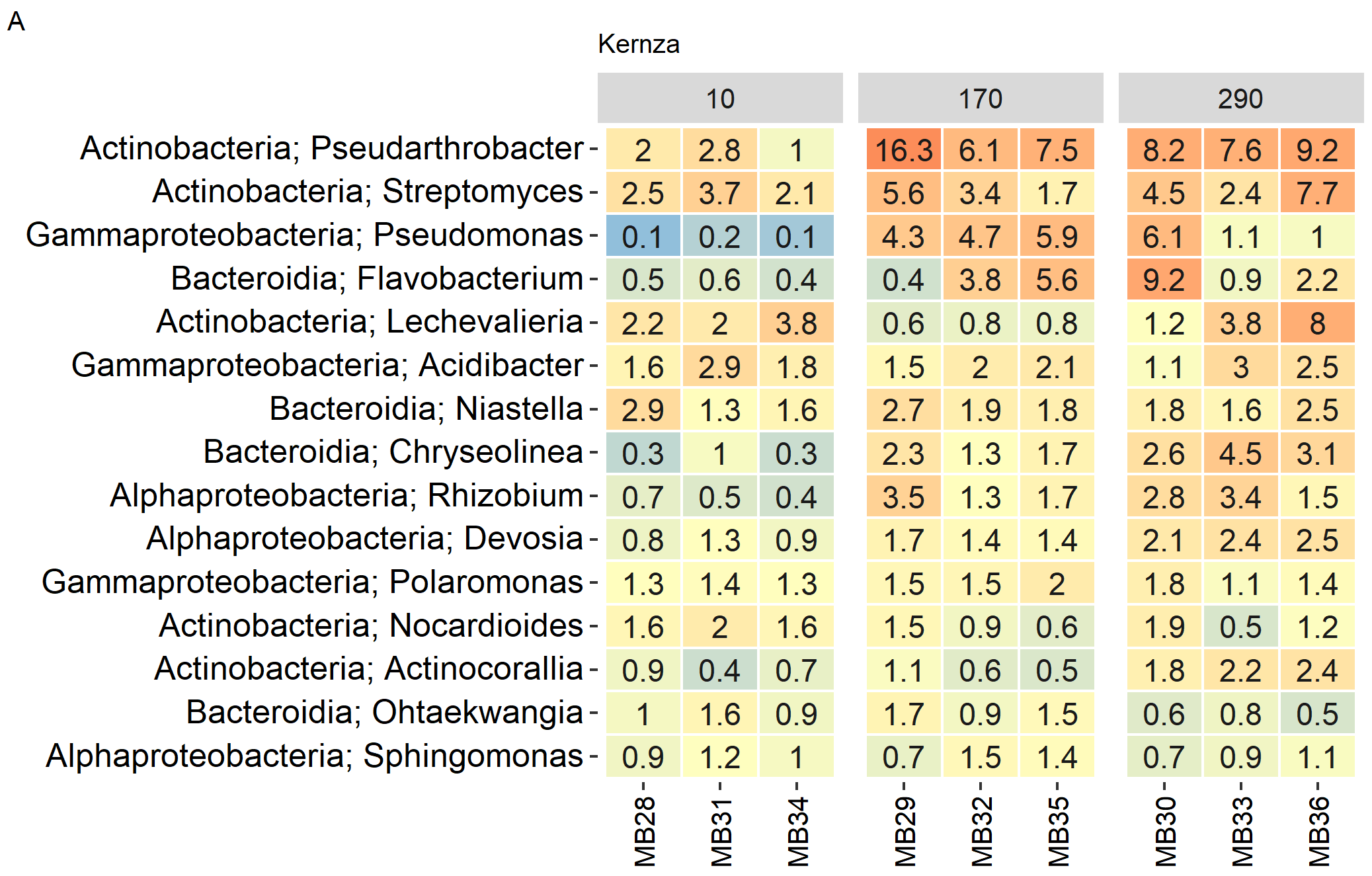

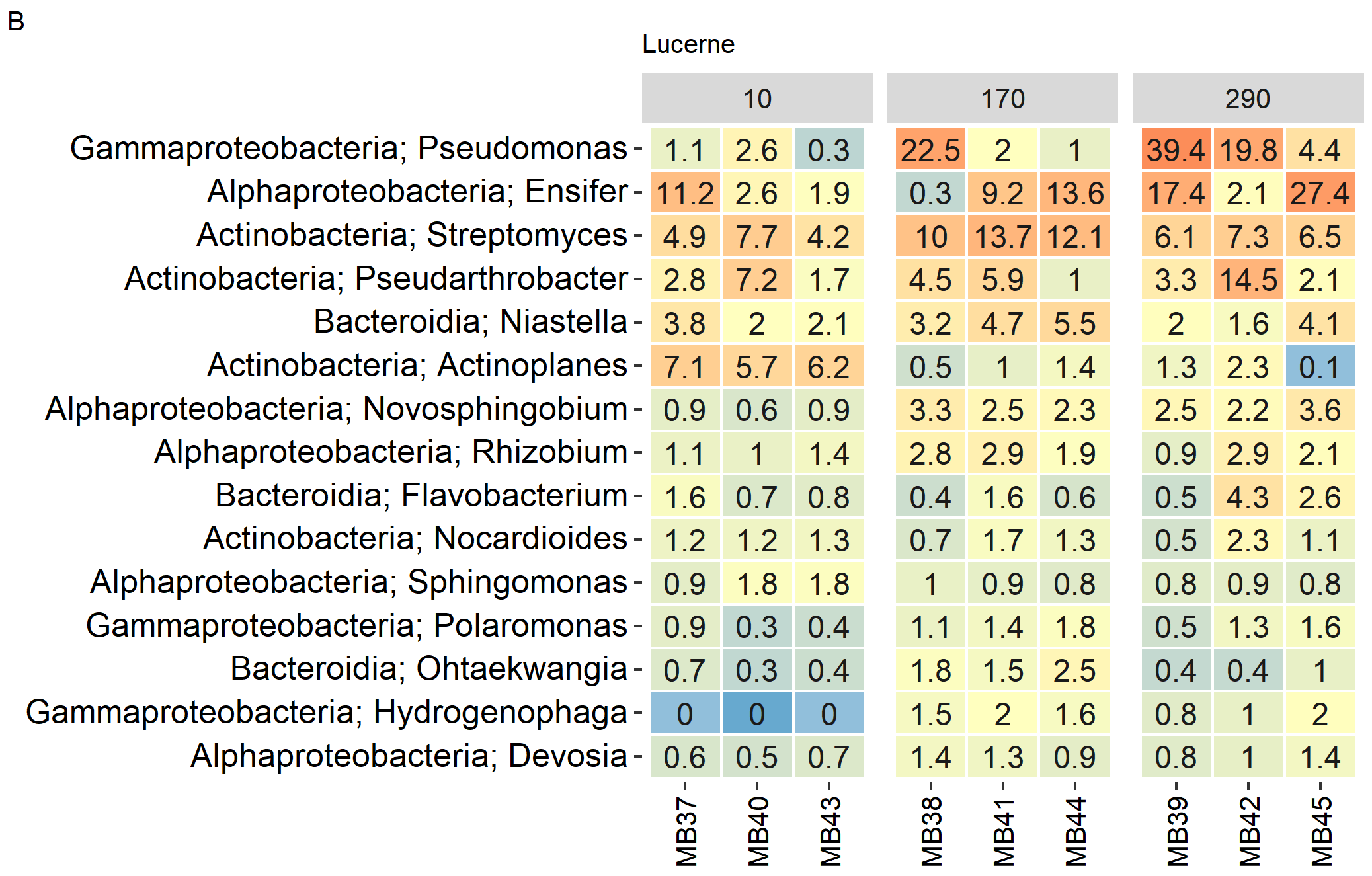

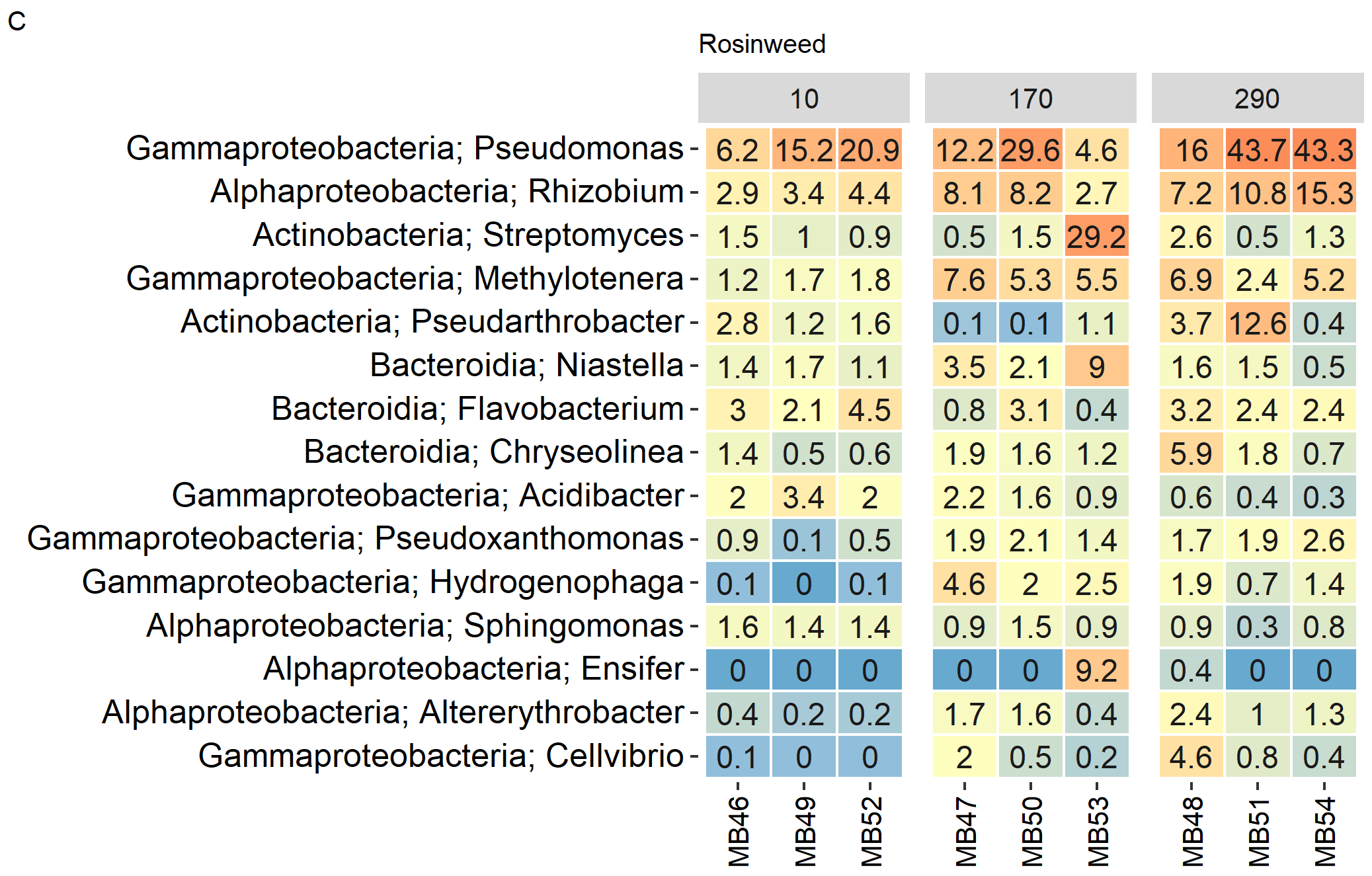

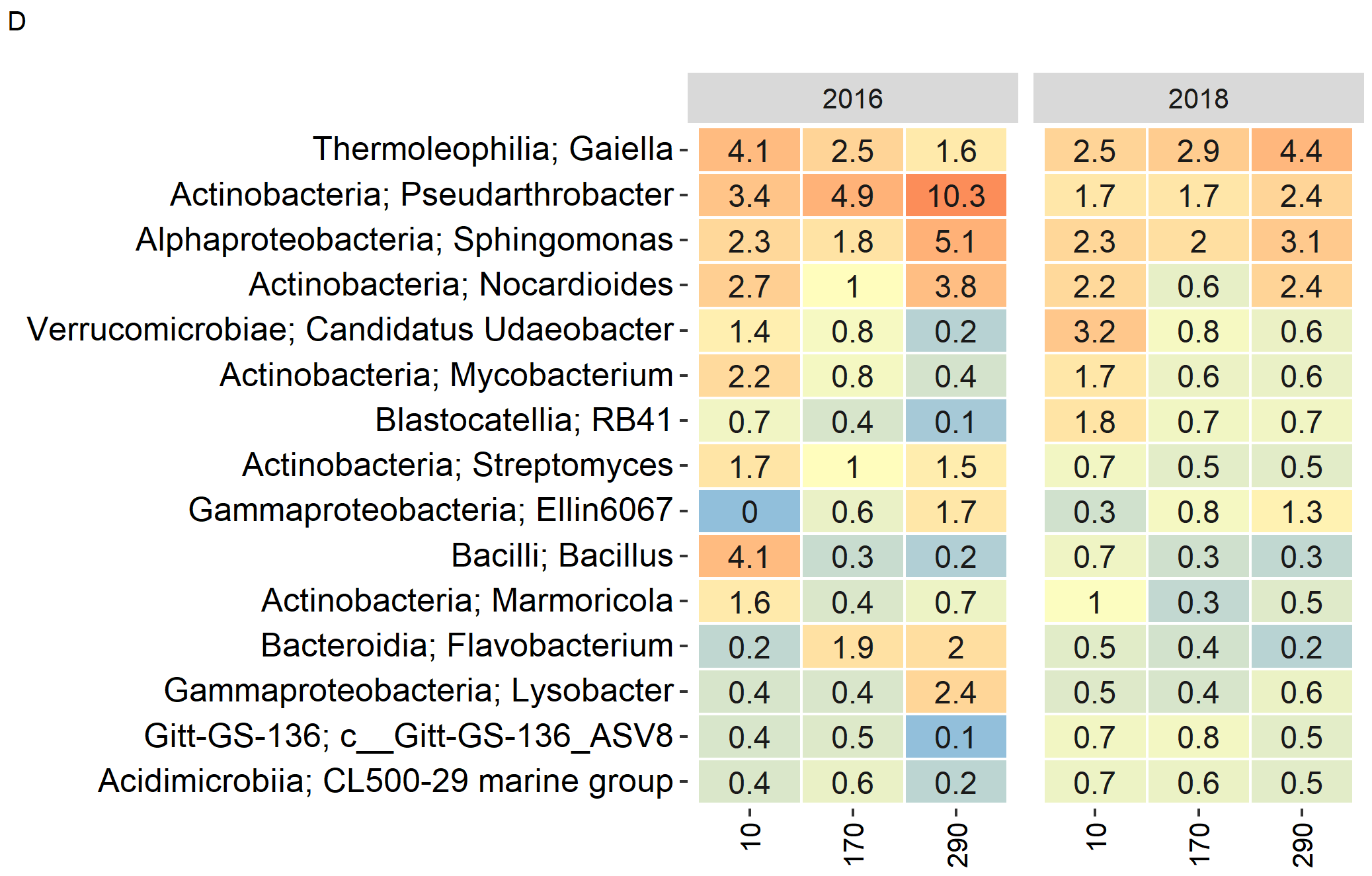

**Supplemental Figure S7**. Venn diagram of the bulk soil samples from all root towers (n = 9). Values indicate number of ASVs and percentages are the proportion of ASVs out of all ASVs in the bulk soil samples.

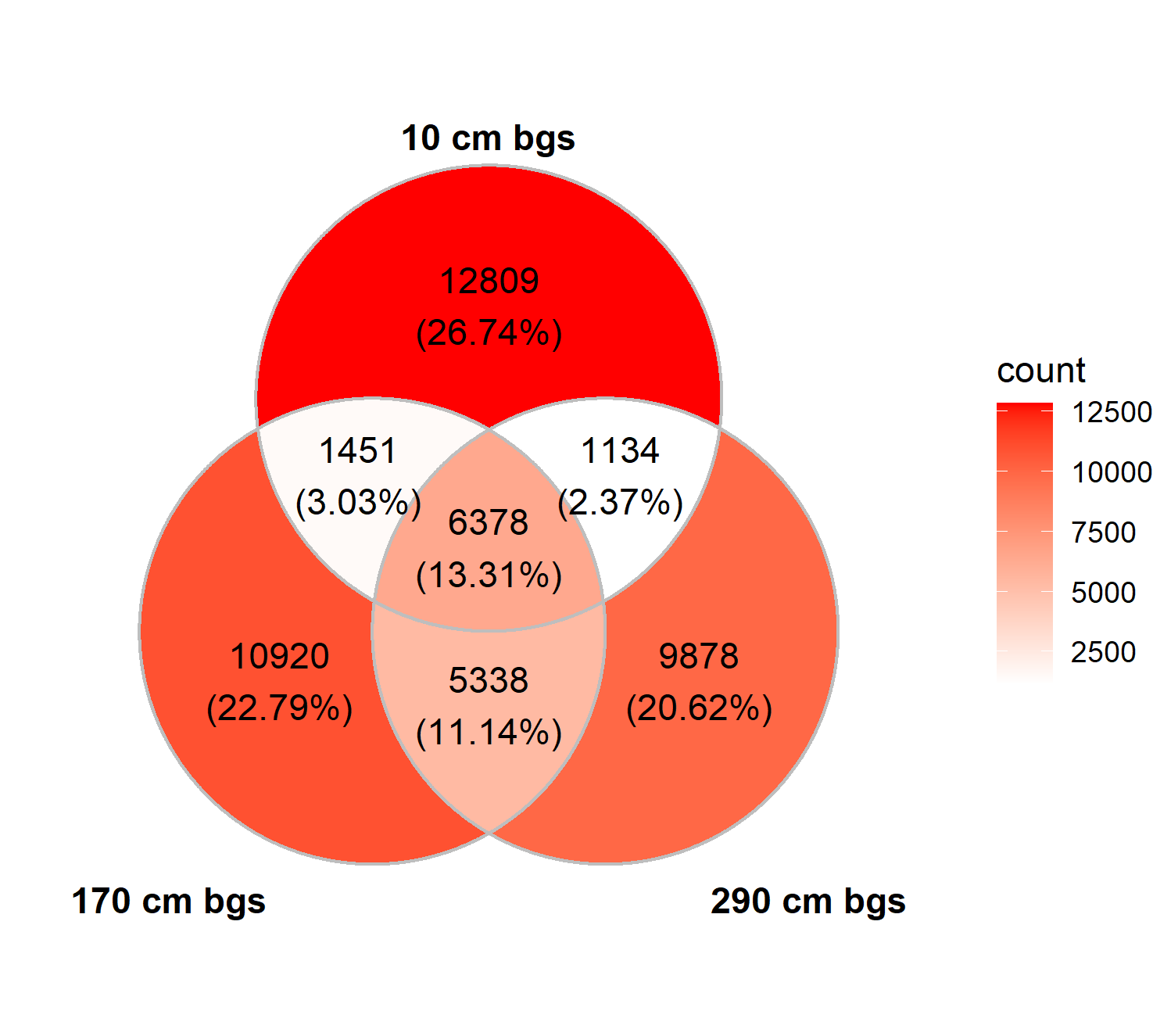

**Table S1**. Primer sequences, annealing temperatures and reaction efficiencies for the qPCR reactions used in this study.

| **Target region** | **Primers** | **Sequences (5’ to 3’)** | **Annealing Temperature** | **Reference** | **Reaction Efficiency** |
| --- | --- | --- | --- | --- | --- |
| 16S rRNA | 341F | CCT AYG GGR BGC ASC AG | 58°C | (Yu *et al.* 2005) | 95.8-96.5% |
|  | 805R | GAC TAC HVG GGT ATC TAA TCC |  | (Herlemann *et al.* 2011) |  |
| ITS 1 | ITS1F | CTT GGT CAT TTA GAG GAA GTA A | 56°C | (Gardes and Bruns 1993) | 83.3-86.1% |
|  | ITS2 | GCT GCG TTC TTC ATC GAT GC |  | (White *et al.* 1990) |  |
| nirK | nirK-F1aCu | ATC ATG GTS CTG CCG CG | 58°C | (Hallin and Lindgren 1999) | 88.8 -99.5% |
|  | nirK-R3Cu | GCC TCG ATC AGR TTG TGG TT |  | (Hallin and Lindgren 1999) |  |
| nosZ | nosZ-F | CGY TGT TCM TCG ACA GCC AG | 63°C | (Kloos *et al.* 2001) | 94.8-96.5% |
|  | nosZ-1622R | CGS ACC TTS TTG CCS TYG CG |  | (Throbäck *et al.* 2004) |  |
| nifH | nifHF | AAA GGY GGW ATC GGY AAR TCC ACC AC | 55°C | (Rösch and Bothe 2005) | 82.5% |
|  | nifHRb | TGS GCY TTG TCY TCR CGG ATB GGC AT |  | (Rösch and Bothe 2005) |  |
| nirS-1 | nirS cd3AF | GTS AAC GTS AAG GAR ACS GG | 55°C | (Throbäck *et al.* 2004) | 79.8-82.1% |
|  | nirS R3cd | GAS TTC GGR TGS GTC TTG A |  | (Throbäck *et al.* 2004) |  |
| amoA | amoA-1Fmod | CTG GGG TTT CTA CTG GTG GTC | 58°C | (Meinhardt *et al.* 2015) | 102.6-106.1% |
|  | GenAOBR-1 | GCA GTG ATC ATC CAG TTG CG |  | (Meinhardt *et al.* 2015) |  |

**Table S2.** PERMANOVA results for 16S rRNA amplicon data using plant species, depth and the interactions as factors. The analysis was based on Bray-Curtis dissimilarity index and 1,000 permutations.

|  |  | Rhizosphere | | | |  | Bulk Soil | | | |
| --- | --- | --- | --- | --- | --- | --- | --- | --- | --- | --- |
| Factor | Df | SumOfSqs | R^2^ | F | P |  | SumOfSqs | R^2^ | F | P |
| Depth | 2 | 1.808 | 0.254 | 7.313 | 0.001 |  | 3.305 | 0.649 | 24.42 | 0.001 |
| Plant Species | 2 | 2.015 | 0.283 | 8.152 | 0.001 |  | 0.220 | 0.043 | 1.626 | 0.115 |
| Depth:Plant Species | 4 | 1.062 | 0.149 | 2.149 | 0.001 |  | 0.350 | 0.069 | 1.293 | 0.199 |
| Residual | 18 | 2.225 | 0.313 |  |  |  | 1.218 | 0.239 |  |  |
| Total | 26 | 7.109 | 1.00000 |  |  |  | 5.093 | 1.00000 |  |  |

**Table S3**. The mean Bray-Curtis dissimilarity within (3 comparisons) or between species (9 comparisons) for each depth. A value of 1 means that no Species are shared between two communities, 0 means that two communities are identical.

| Depth [cm bgs] | Lucerne | Kernza | Rosinweed | Lucerne/ Kernza | Lucerne/ Rosinweed | Kernza/  Rosinweed |
| --- | --- | --- | --- | --- | --- | --- |
| 10 | 0.42 | 0.49 | 0.47 | 0.63 | 0.71 | 0.74 |
| 170 | 0.42 | 0.52 | 0.59 | 0.62 | 0.65 | 0.72 |
| 290 | 0.53 | 0.55 | 0.52 | 0.72 | 0.78 | 0.79 |
